# Integration of mass-spectrometry-based global metabolomics and proteomics analysis to characterise different senescence induced molecular sub-phenotypes

**DOI:** 10.1101/2022.11.30.518588

**Authors:** Domenica Berardi, Gillian Farrell, Abdullah Alsuntan, Ashley McCulloch, Zahra Rattray, Nicholas JW Rattray

## Abstract

Cellular senescence is a key driver of ageing and its related disease. Thus, targeting and eliminating senescent cells is a major focus in biogerontology to predict and ameliorate age-related malady. Many studies have focused on targeting senescence through the identification of its molecular biomarkers. However, these are not specific for senescence and have different expression patterns across various senescence phenotypes. Here we report a combination of molecular studies (ß-galactosidase expression, DNA damage and replication immunodetection) with a mass spectrometry analysis integrating intra and extracellular global metabolomics to reveal small molecules differentially expressed across multiple senescence phenotypes (replicative senescence, x-ray, and chemical-induced senescence).

Altered key intracellular metabolic changes were identified, depending on the stress stimuli, which were consistent with the presence of pro-inflammatory metabolites in the cellular secretome.

Our work shows the advantage of combining molecular and metabolomics studies for the detailed analysis of cellular senescence and that senescence phenotype changes upon induction method.

## INTRODUCTION

Currently, more than 12% (900 million) of the global population are over 60 years old and by 2050 this estimate will raise up to 22% (2 billion) [1]. This demographic change will lead to increased medical care and social needs, therefore having a radical impact on the structure and function of society, the global economy and health systems [2].

Human ageing is a temporal process of structural and functional accruement of damage at the molecular, cellular and tissue level which leads to the state of disease development, that can turn into a chronic condition and ultimately leading to death [3]. Accordingly, ageing represents the primary risk factor in the majority of diseases like diabetes, cancer [4], neurological [5], cardiovascular, and immune disorders [6]. For this reason, strategies to improve human health span, through the prevention and amelioration of age-related disease, have been extensively studied [7]. A better understanding of the biology of ageing is needed to develop diagnostics and therapies capable of tracking and targeting features of ageing.

Among the primary feature of ageing there is the accumulation of senescent cells, which have been observed in several age-related diseases and in the elderly [8]. Senescence is a hallmark of ageing characterised by the arrest of cell proliferation and resistance to death [9]. At present, induction of the senescence phenotype in normal human cells is the widest accepted strategy in ageing research to better understand its biological mechanisms and role in pathological conditions [10–12].

Cellular senescence was first reported in normal human fibroblasts where replication-associated telomere shortening was observed, which has widely been used as a prototype model for senescence known as replicative senescence [13, 14]. Stress-induced premature senescence (SIPS) is another type of senescence which includes DNA damage-induced senescence, Oncogene induced senescence, mitochondrial dysfunction-induced senescence, epigenetically induced senescence, and reactive oxygen species (ROS) induced senescence [15–17].

Several SIPS have been reported that differ based on the expression of specific senescence features, which include DNA damage, telomere attrition, cell cycle arrest, resistance to apoptosis, endoplasmic reticulum stress, increased lysosomal content of ß-galactosidase (ß-Gal), mitochondrial dysfunction, senescence-associated heterochromatin foci (SAHF), flattened morphology, larger cell surface, senescence-associated secretory phenotype (SASP), and metabolic reprogramming [17, 18] However, these features are facultative in the senescence phenotype and dependent on senescence-inducing stimuli and the cell type impacted.

Ionizing radiation (including X-rays) induce DNA damage and mutations directly by secondary electrons emission and/or indirectly by ROS formation [19], therefore resulting in DNA breaks (single or double stranded), DNA-protein crosslinks, oxidation of bases, and formation of abasic sites [20]. Hydroxyurea is an inhibitor of ribonucleotide reductase [21] by removing tyrosine free radicals that are required for the reduction of nucleoside diphosphates and DNA synthesis, consequently resulting in DNA single -strand breaks [22]. Etoposide is a chemotherapy agent that inhibits DNA topoisomerase II (TopII) therefore preventing the re-ligation of broken DNA strands [23].

The heterogeneity of the senescence phenotype and lack of sensitive selective markers of senescence pose a significant challenge for the identification of senescent cells in culture, tissues and *in vivo* [24]. Combining the measurement of multiple molecular biomarkers is the current strategy used for the identification of senescent cells.

Metabolomics is the comprehensive analysis of small molecule metabolites, which are downstream by-products of gene, RNA, and protein function [25]. Metabolomics can provide an integrated portrayal of biological pathways and how changes in biochemistry are associated with human disorders and disease. Considering that the application of metabolomics to ageing research is relatively new, it is still lacking an exhaustive annotated metabolic dataset describing the complexity of senescence phenotypes.

Here we report the study of molecular and metabolic markers of different senescence phenotypes through measuring the expression of ß-Gal, DNA damage foci, cell cycle activity, morphology, and global intracellular and extracellular metabolites to analyse changes in the phenotype of senescent cells in response to different methods of induction in normal human fibroblasts.

## RESULTS

### Induction of senescence in normal human fibroblasts

The initial focus of this work was to design and classify the biomolecular differences of functional models of replicative and stress-induced senescence. To achieve this, we passaged cells multiple times in culture and treated them with different doses of X-rays, hydroxyurea and etoposide as previously outlined (see methods). The state of senescence was optimised for each condition with a target cell viability of ~ 50%, > 50% β-galactosidase (β-Gal) expression in sub-optimal conditions (pH<6) and an increased surface area of treated cells compared to their relative controls. The rationale behind this optimisation, was to ensure an adequate number of cells (biomass) for subsequent imaging and metabolomics experiments.

**Figure 1.**
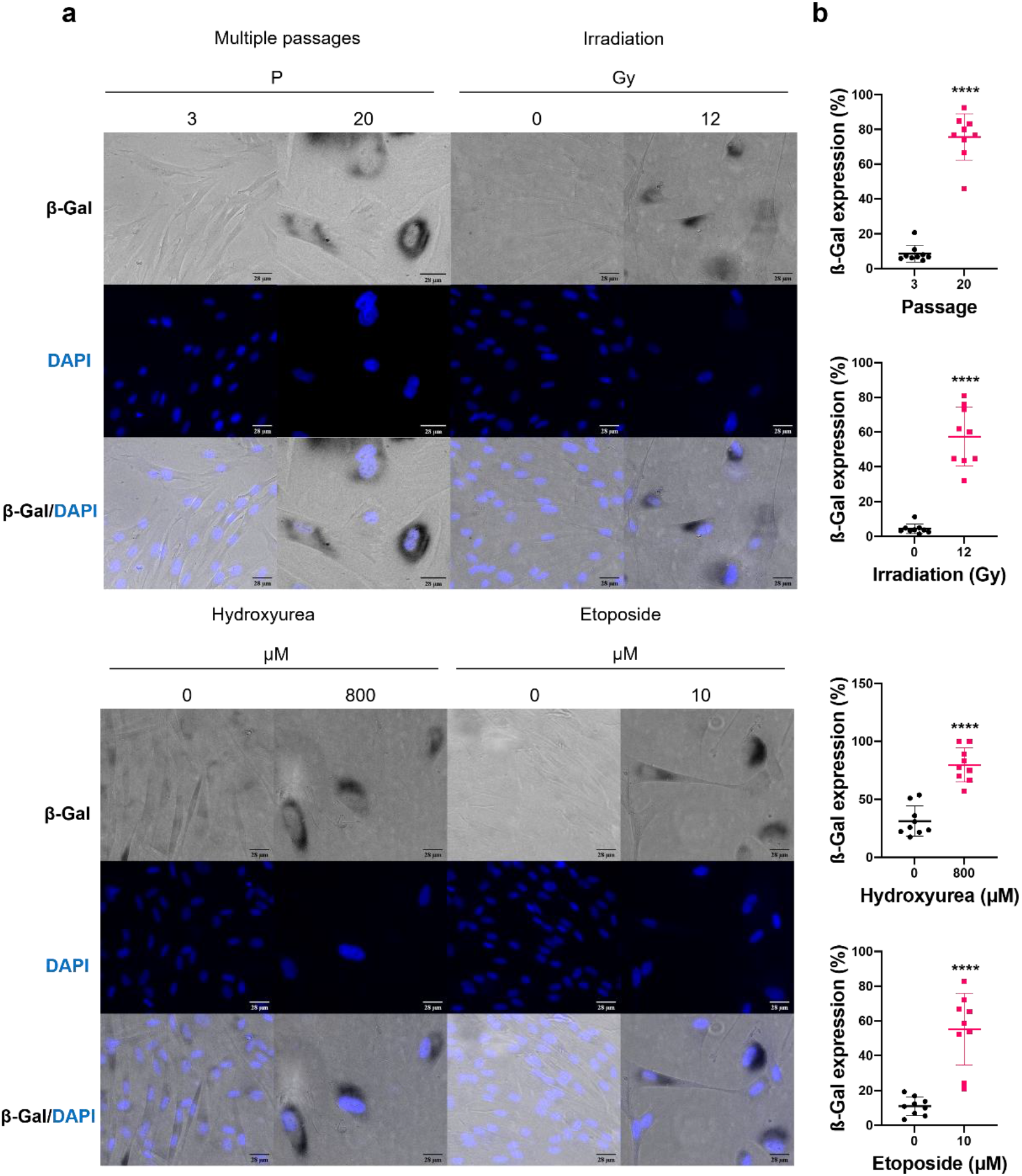
β-Gal in senescence-induced HFF-1 cells. Representative phase contrast images of β-Gal staining (grey), DAPI (blue) and composite (β-Gal (grey) and DAPI (blue)) in cells at passage 20,1 week post irradiation at 12 Gy, 800μM hydroxyurea for two weeks and 10μM etoposide for one week (a). Corresponding β-Gal levels expressed as a percentage of cell count relative to the number of nuclei counted using ImageJ (b). 9 repeats from 3 independent replicates with on average >50 cells per each sample. p-values have been determined through t-test and represented as ≤0.00005=****.

Based on our results, senescence was induced across all the conditions examined at a late passage (P=20) and higher ionizing radiation dose and chemical treatment compared to control. We observed ~ 80% of cells were β-Gal-positive after 20 passages and treatment with 800 μM hydroxyurea, while cells treated with 12 Gy IR and 10μM etoposide resulted in >60% β-Gal-positive cells. Of notice, the control samples for each condition (passage 3, 0 Gy IR, and 0 μM of hydroxyurea and etoposide) had a baseline percentage of β-Gal-positive cells which corresponds to <20% in the passaged, irradiated and etoposide-treated cells after a one week incubation, while ~30% in the hydroxyurea-treated cells following a two week incubation. These results show a baseline number of senescent cells exist under normal cell culture conditions and that β-Gal expression is influenced also by the experiment incubation time.

### Senescence results in the formation of γH2AX and Ki-67 foci

Senescent cells are characterised by accumulation of DNA damage and cell cycle arrest [26]. Therefore, we next analysed the extent to which late passages in culture (P=20), exposure to 12Gy IR, 800μM hydroxyurea and 10μM etoposide promotes the accumulation of DNA double-strand breaks (DSBs) and cell cycle arrest. Phosphorylated histone H2 variant γH2AX (γH2AX) is a key marker of DNA DSBs [27], while Ki-67-a marker of cellular proliferation-is progressively expressed during the S phase of the cell cycle (replication phase) [28]. To measure the formation of DNA DSBs and cell cycle arrest in senescence-induced cells we performed γH2AX (**Figure 2**).

**Figure 2.**
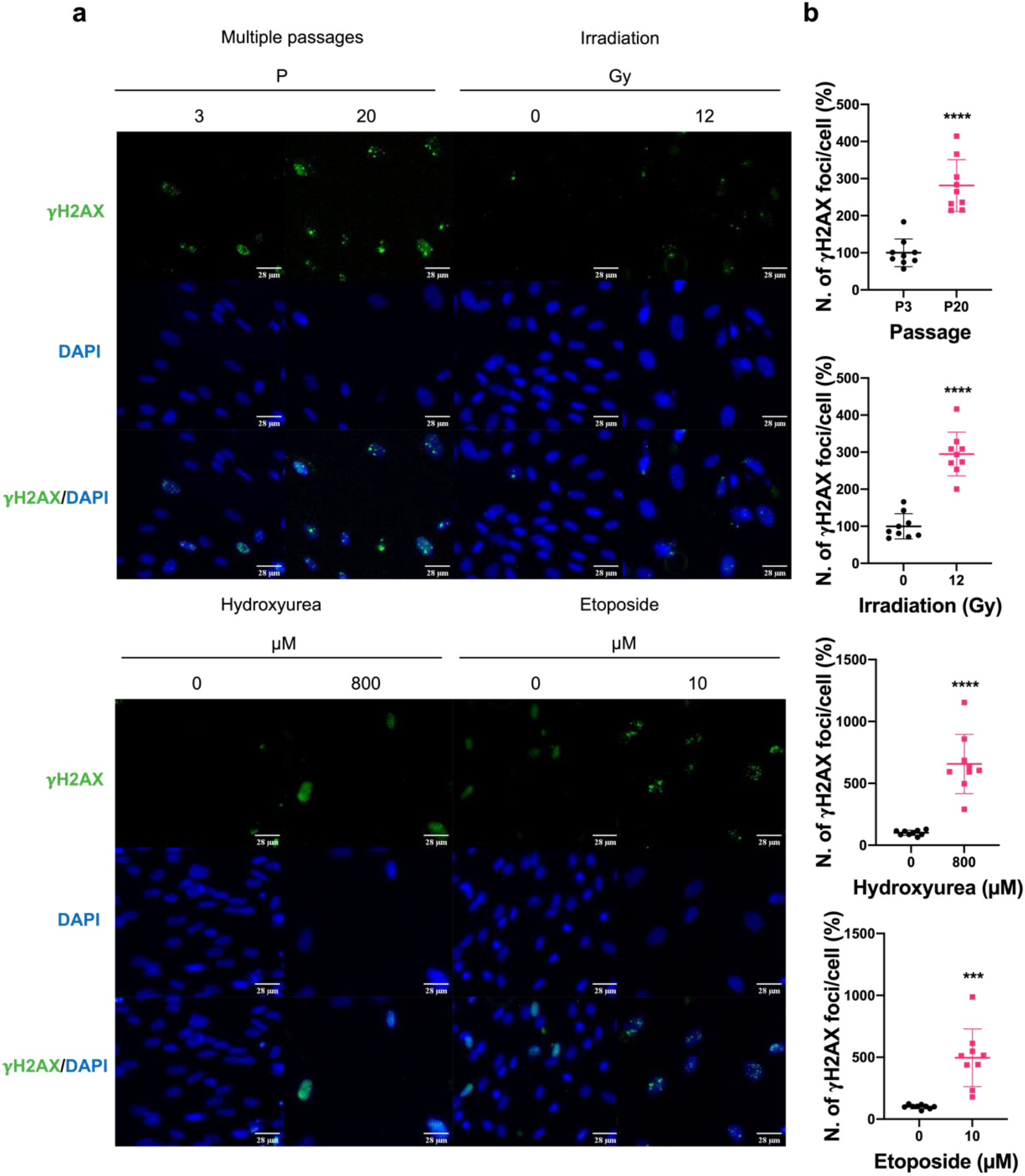
γH2AX foci immunodetection in senescence-induced HFF-1 cells. Representative images of γH2AX immunostaining (green), DAPI (blue) and composite (γH2AX (green) and DAPI (blue)) in cells at passage 20, 12Gy IR for 1 week, 800 μM hydroxyurea for two weeks and 10μM etoposide for one week (a). Corresponding γH2AX foci count per cell as measured in Cell Profiler. 9 repeats from 3 independent replicates with on average >50 cells per each sample. Pairwise comparison was performed, and p-values have been determined using a t-test and represented as: 0.0005=***, <0.00005=****.

Based on our results, γH2AX foci levels increased in HFF-1 cells at later passages (i.e., P=20) and following treatment with 12Gy IR, 800 μM hydroxyurea and 10 μM etoposide **(Figure 2a,b**). Next, cell cycle activity was analysed via Ki-67 immunochemistry (**Figure 3**).

**Figure 3.**
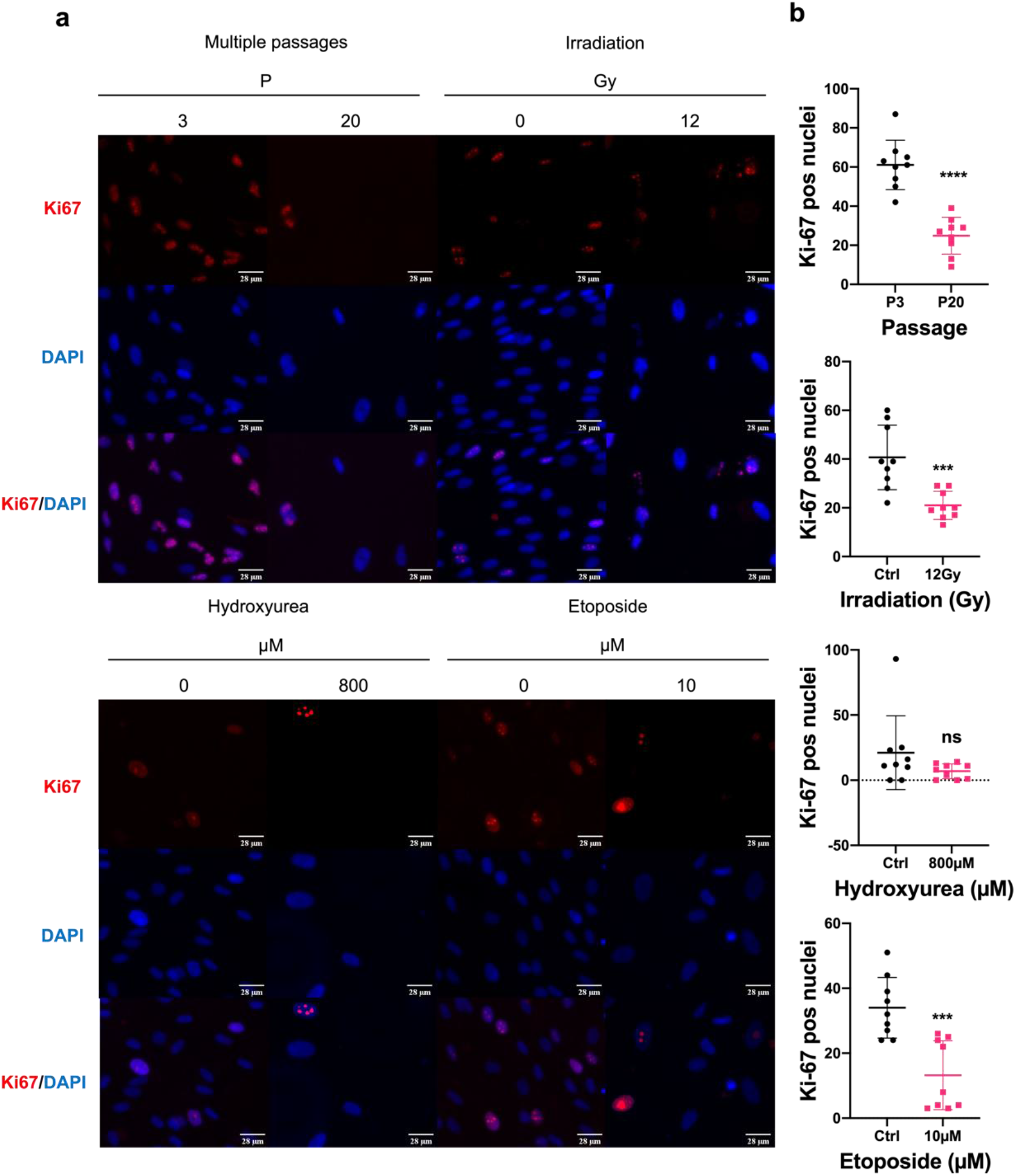
Ki-67 positive nuclei immunodetection in senescence-induced normal human fibroblasts. Representative images of Ki-67 immunostaining (red), DAPI (blue) and composite (Ki-67 (red) and DAPI (blue)) in cells at passage 20, 12Gy IR for one week, 800μM hydroxyurea for two weeks and 10μM etoposide for one week (a). Corresponding Ki-67 expression levels representative of positive nuclei count (b). 9 repeats from 3 independent replicates with on average >50 cells per each sample. p-values have been determined through t-test and represented as: 0.0005=***, <0.00005=****, and non-significant=ns.

Regarding Ki-67 expression, we observed that the number of Ki-67-positive nuclei were reduced in all our samples with a significant reduction in the late passaged, irradiated and etoposide treated cells **(Figure 3a,b)**. An increase of γH2AX foci is consistent with the accumulation of DNA damage in senescence cells both in vitro and in vivo. Reduced levels of Ki-67 foci in the senescence-induced cells reflect their reduced proliferation.

### Senescence-associated metabolic changes vary depending on the method of induction

To investigate the variation among different senescence models, their metabolic content was profiled using the untargeted liquid chromatography-mass spectrometry pipeline illustrated in Appendix 1. Briefly, after data acquisition, data were processed and analysed in Compound Discoverer 3.3. Principal component analysis (PCA) was performed to reduce the global dataset of features from all the conditions, and to cluster and easily visualise the differences between the senescent. Pairwise PCA analysis allowed to better discriminate the differences in the components between controls and treated samples (**Figure 4a**). Differential analysis through volcano plot provided information on the differential number of significant metabolites altered (enriched or depleted) upon induction of senescence through multiple passages, ionizing radiation, and incubation with drugs (i.e., hydroxyurea, and etoposide) (**Figure 4**).

**Figure 4.**
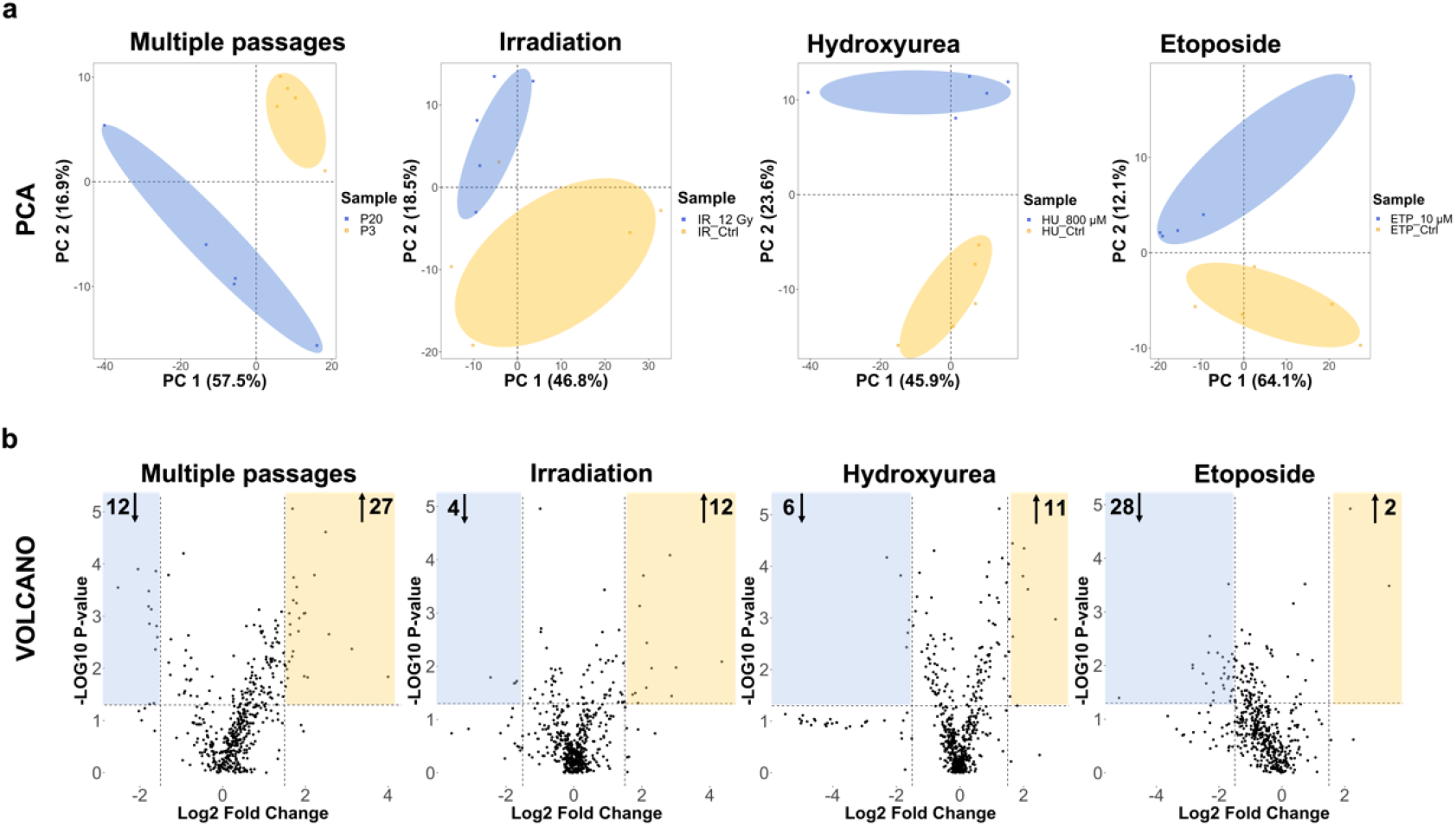
Statistical analysis of the metabolic features identified in HFF-1 cells. Cells at passage 20, and following 12Gy IR for one week, 800μM hydroxyurea for two weeks and 10 μM etoposide for one week. A) PCA pairwise analysis of metabolites altered in the different senescence-induced cells. For each treatment group, five replicates were used. Data points in the two-dimensional PCA score plot were central scaled. B) volcano plots displaying enriched (yellow) and depleted (blue) metabolic features by representing the log2 fold change in altered features and the -log10 adjusted p-values with cut off values selected at >1.5 and <0.05, respectively.

Pooled quality control (QC) data confirmed the stability of data acquisition across all the measurements performed in positive/negative switching mode (**Error! Reference source not found.b of Appendix 1**). Feature separation was observed between treated samples and their controls (**Figure 4a**). Volcano plots indicate the differential number of metabolic features that are significantly altered following senescence induction (**Figure 4b**). From our results, we observed that in cells at late passage (p=20) and cells treated with irradiation (12 Gy), and hydroxyurea (800 μM) most of the metabolic features were enriched with respect to their relative control. Conversely, in the etoposide-treated cells (10 μM) the number of depleted metabolic features was higher (n.=28) compared to the enriched features (n.=2) (**Figure 4b**). Of notice, most metabolites were enriched in the passaged, irradiated and hydroxyurea-treated cells, while a higher number of depleted metabolites was detected in etoposide-treated samples. Together, these findings show a differential metabolic response of normal human fibroblasts to different stress stimuli inducing different senescence phenotypes.

### Altered amino acid metabolism in different senescence-induced cells

To analyse the chemical and biomolecular changes in response to the induction of senescence, we used MetaboAnalyst to identify specific metabolic pathways altered at later passages in culture (P=20), following 12Gy IR, and treatment with 800μM hydroxyurea and 10μM etoposide. Thus, we performed pathway analysis for all the conditions (**Figure 5**) by integrating the list of identified metabolites with the list of senescence-associated genes selected from the “The Ageing Gene Database” (GenAge). Among the pathways ranked in the top 10, we selected altered pathways with a corresponding pathway impact >0.1, and FDR p-value <0.05 (**Error! Reference source not found. of Appendix 1**).

**Figure 5.**
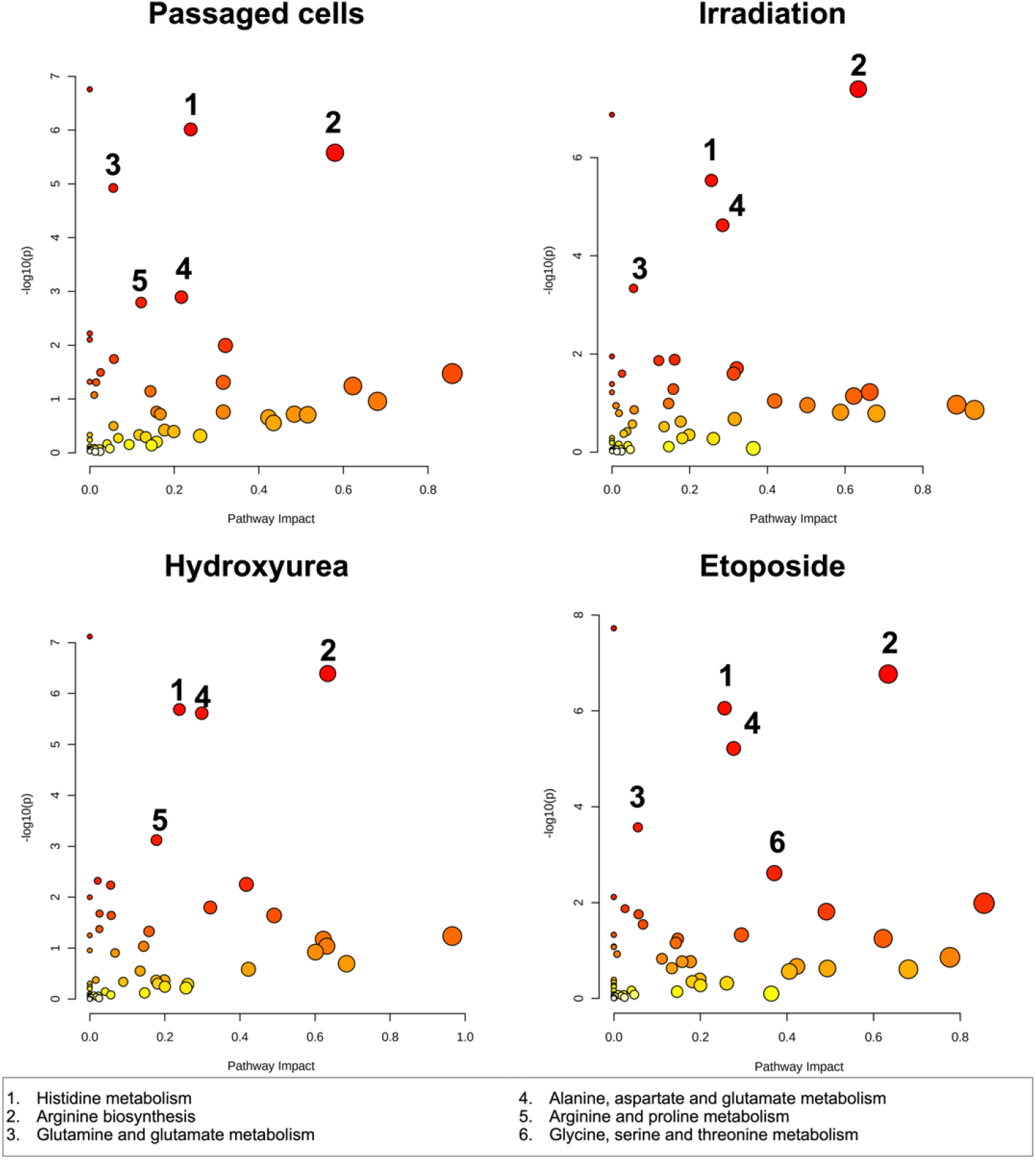
Pathway enrichment analysis of HFF-1 cells. Cells at passage 20 in culture, following 12Gy IR for one week, 800μM hydroxyurea for two weeks and 10μM etoposide for one week. Numbering for each metabolic pathway is kept consistent across the different plots. Pathway analysis was based on the hypergeometric test. Topological analysis was based on betweenness centrality. The tight integration method was used by combining genes and metabolites into a single query. A FDR p<0.05, and pathway impact >0.1 were deemed significant.

Across the senescence model examined, the top putative pathways significantly altered in MetaboAnalyst were based on amino acid metabolism (histidine, arginine, alanine, aspartate, glutamate and lysine) for replicative and irradiation-induced senescence; and lipid metabolism (pantothenate and CoA biosynthesis, and glycerophospholipid metabolism) for hydroxyurea and etoposide-treated cells. Following the identification of metabolic pathways altered upon senescence induction, we constructed a Venn to outline common altered metabolic features.

Overlapping pathways are represented by histidine metabolism, arginine biosynthesis, alanine, aspartate and glutamate metabolism, suggesting that the senescent state is associated with changes in the metabolism of these amino acids. Arginine and proline degradation pathways are common to late passaged and hydroxyurea-treated cells. Changes in glutamine and glutamate metabolism are relevant for late passaged and irradiated cells. Glycine, serine, and threonine metabolism are specifically relevant to etoposide-treated cells.

Next, we evaluated the relative changes in the levels of the individual metabolites that are representative of the altered metabolic pathways in the different senescence-induced models. The results were presented through a heatmap clustering analysis (**Figure 6**). A wider list of compounds specific to each model of senescence is provided in **Appendix 1**.

**Figure 6.**
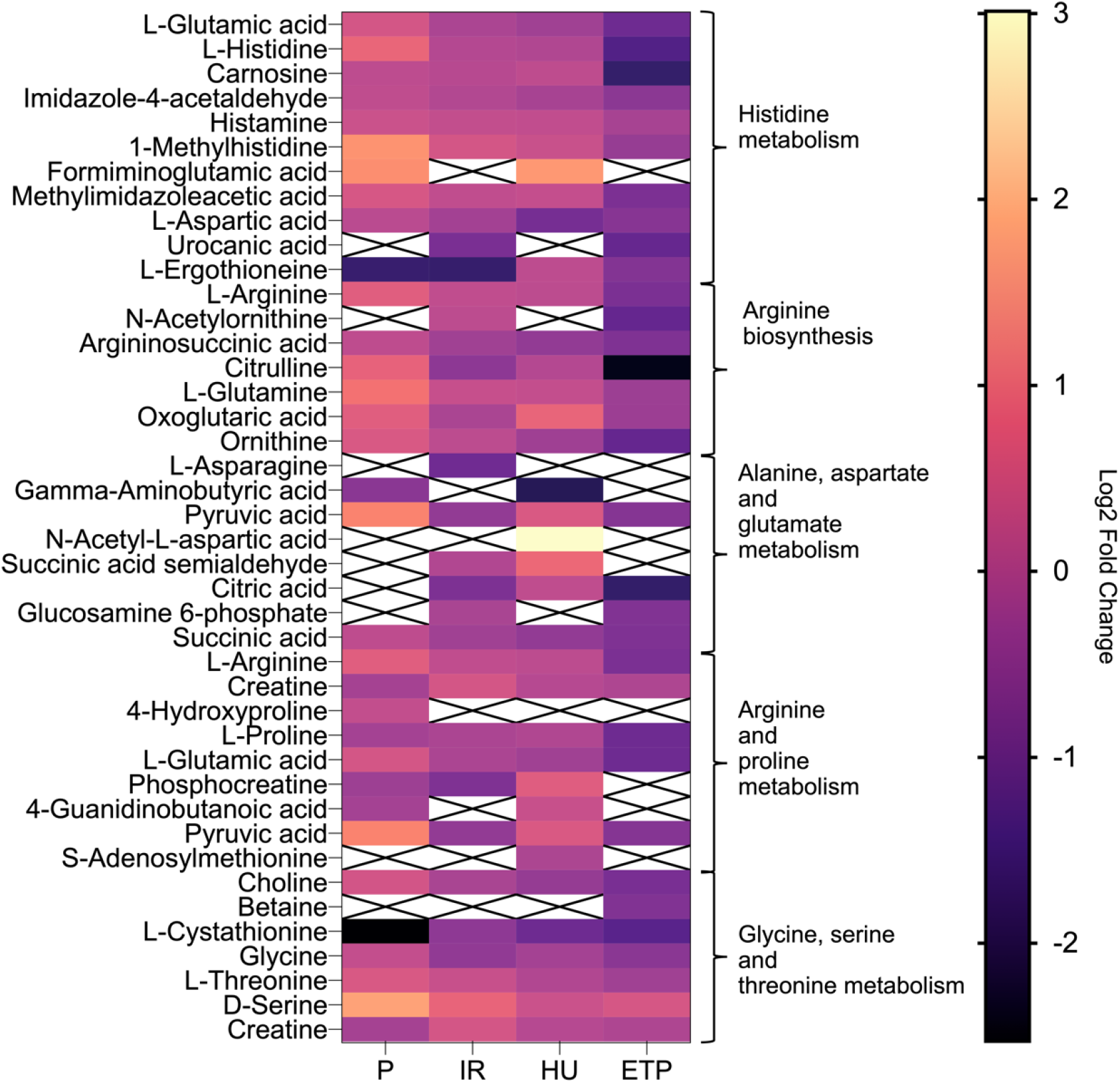
Heatmap cluster analysis. Study of relevant metabolites associated with the pathways altered upon the induction of senescence through multiple passages (P=20), irradiation (12Gy), hydroxyurea (800μM) and etoposide (10μM). Clustering reflects the relevant enriched metabolic pathways including histidine metabolism, arginine biosynthesis, alanine, spartate and glutamate metabolism, glycine, serine and threonine metabolism. X boxes indicate the absence of detected peak for the metabolite of interest.

Analysis of the heatmap cluster (**Figure 6**) indicates significantly high levels of serine and hypoxanthine were detected in cells at late passage in culture and following treatment with 12Gy IR. High levels of serine were detected also in the hydroxyurea and etoposide-treated cells although the enrichment was not significant relative to the control. Significant depletion of taurine and hypotaurine was a commonly observed feature of all the senescent cells in this study, accompanied by a high ratio of GSSG/GSH calculated for replicative senescent cells (0.6), hydroxyurea (1.3) and etoposide-treated cells (1.3).

Enrichment of alpha-ketoglutarate (α-KG), methionine, 1-methylhistidine and carnosine was a common feature of late passaged cells, and cells treated with ionizing radiation and hydroxyurea. However, this feature was absent in cells treated with etoposide. Depletion of proline, glutamate and aspartate have been observed in all the senescence-induced cells although not at significant levels. Significantly depleted levels of acetylcholine have been shown in cells treated with hydroxyurea and etoposide, except in cells at late passage.

Taken together, these results highlight the prevalence of depleted metabolites upon treatment with etoposide compared to the passaged, irradiated and hydroxyurea-treated cells.

An overview of the metabolic features altered in response to senescence induction is provided in **Figure 7**, where we mapped the differences in metabolite levels of the various senescence models using the Kyoto Encyclopaedia of Genes and Genomes (KEGG) database.

**Figure 7.**
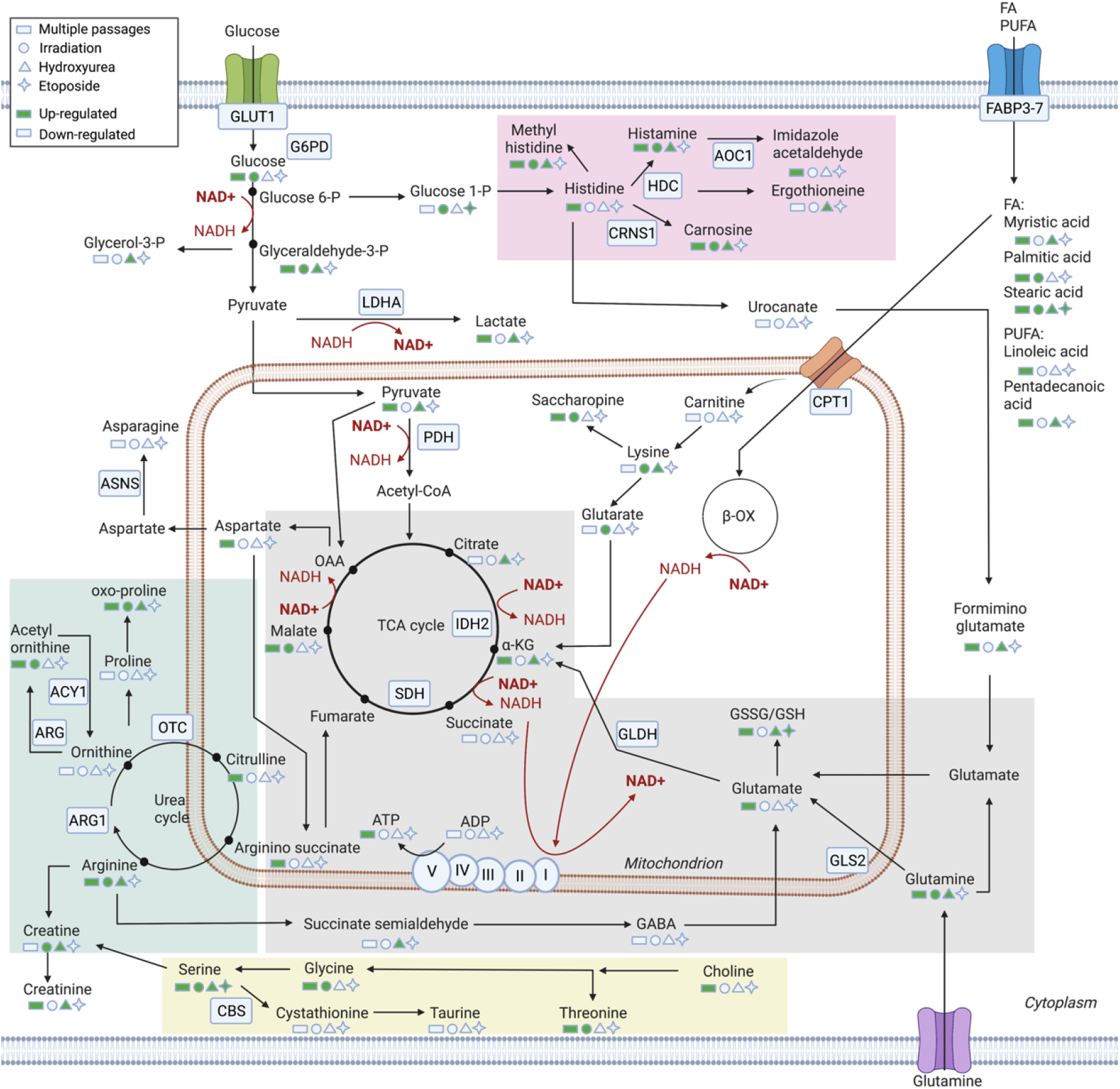
Summary of putatively identified metabolic pathways altered upon induction of senescence. Enriched metabolic pathways are represented in the coloured boxes: histidine (red), arginine and proline (green), alanine, aspartate, glutamate and glutamine (grey), glycine, serine and threonine metabolism (yellow).

The enzymatic genes associated with the metabolic reactions have been represented in the map, followed by network analysis with relevant senescence genes. We distinguished among the metabolic changes occurring for each different senescence-induced phenotype. In the replicative senescence model we observed the enrichment of glucose and glutamine metabolism paralleled by enriched TCA acids and metabolites involved in the urea cycle, serine and histidine metabolism. Similarly, in the senescence model induced through irradiation, enrichment of glucose and glutamine was detected, however the TCA acids were mostly depleted. Serine and histidine derived metabolites were also enriched in the irradiated cells. Regarding hydroxyurea-induced senescence, we identified enrichment of glutamine, but depleted levels of glucose. High levels of TCA acids and histidine-derived metabolites were present in the hydroxyurea-treated cells, together with enhanced expression of oxidised glutathione. Finally, in the etoposide-induced senescence phenotype, most of the metabolic pathways were depleted but we measured enrichment of serine and saturated fatty acids similarly to the other senescence phenotypes.

Overall, through the joint metabolic study of the different senescence-induced models we computed similarities and differences in their intracellular metabolic response to different stress stimuli.

### Detection of extracellular inflammatory metabolites in different senescence-induced phenotypes

To analyse the extracellular composition of metabolites, we next applied the metabolomics pipeline to the study of growth media collected from all the senescence-induced cells of this study. We used PCA analysis and volcano plots to examine the differences between growth media aspirated from senescence-induced cells and their relative controls (**Figure 8**).

**Figure 8.**
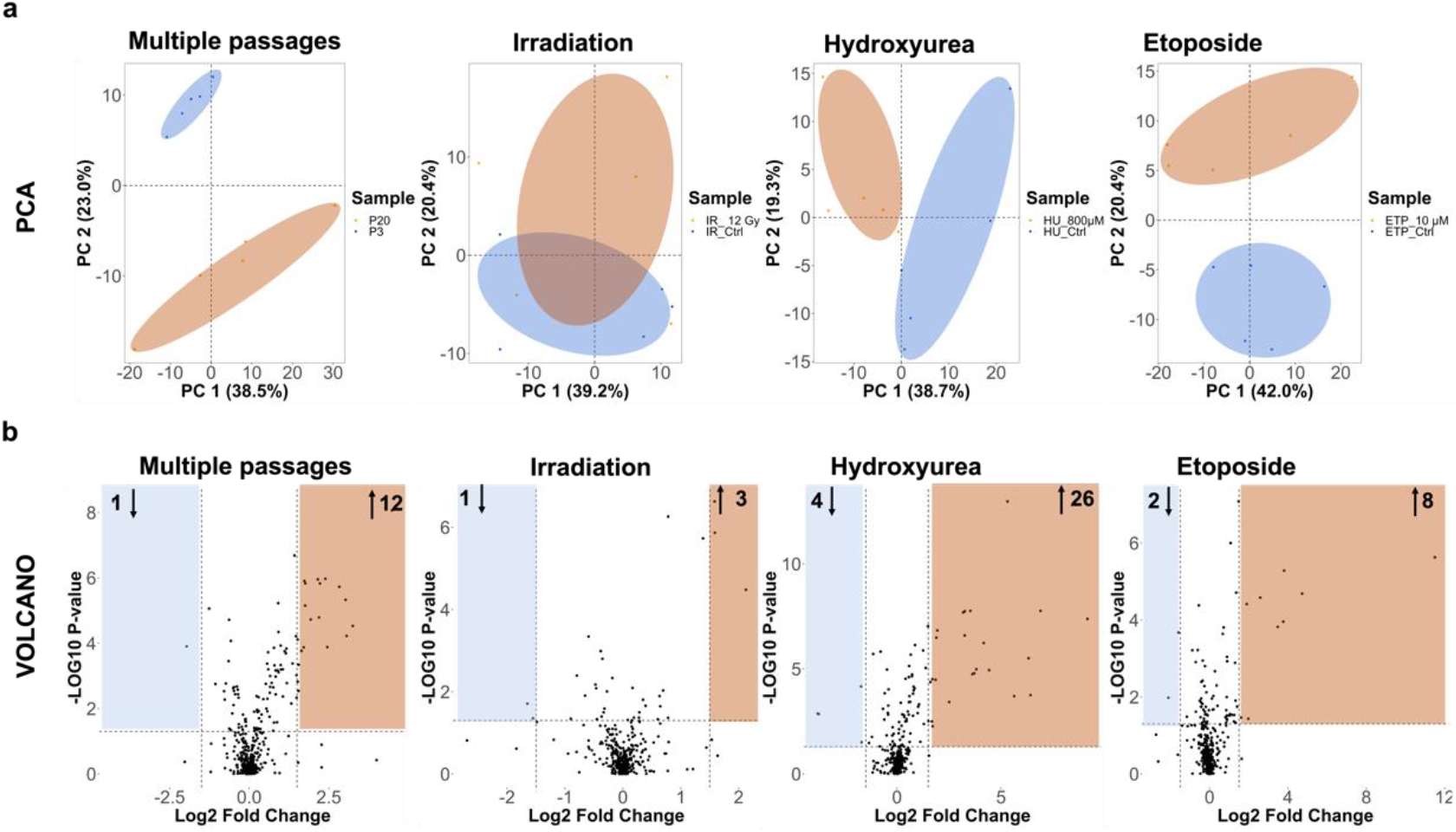
Statistical analysis of the metabolic features identified in the growth media of HFF-1 cells. Cells at passage 20, 12 Gy irradiation for 1 week, 800 μM hydroxyurea for 2 weeks and 10 μM etoposide for 1 week. A) PCA pairwise analysis of metabolites altered in the different senescence-induced cells. For each treatment group, five replicates were used. Data points in the two-dimensional PCA score plot were central scaled. B) volcano plots displaying enriched (yellow) and depleted (blue) metabolic features by representing the log2 fold change in altered features and the -log10 adjusted p-values with cut off values selected at >1.5 and <0.05, respectively.

LC-MS stability was confirmed by the analysis of QC samples. Grouping of different features was observed across all the conditions (**Figure 8a**). Using volcano plots we analysed the differential expression between the metabolites significantly altered in the senescence-induced cells (enrichment and depletion) relative to their controls (**Figure 8b**).

Next, we performed enrichment analysis through MetaboAnalyst with the global list of metabolites and ranked them based on their class. Through a Venn diagram we presented the differences between the enriched metabolites within the growth media and cells of the late passaged cells and those treated with IR, hydroxyurea and etoposide (**Figure 9**).

**Figure 9.**
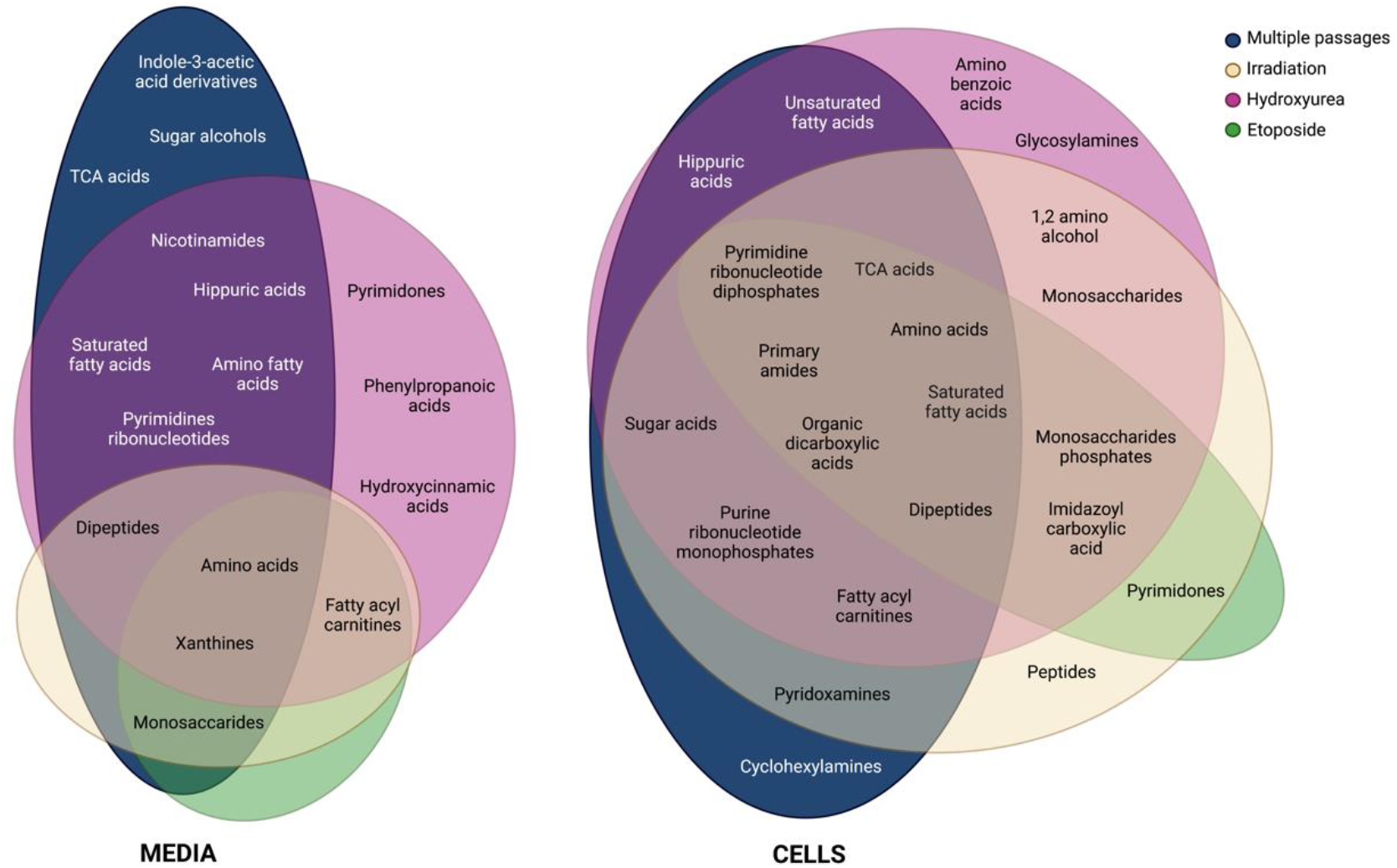
Venn diagram. The diagram represents the enriched class of extracellular and intracellular metabolites in late passaged, irradiated, hydrozyurea and etoposide treated cells.Data have been processed in MetaboAnalyst 5.0.

From analysis of metabolic features altered in the growth media of senescence-induced models (passage, IR and chemically-induced) we observed that amino acids and xanthines represented the most commonly enriched class of metabolite across all senescence-induced models. TCA acids, sugar alcohols and indole-acetic acid derivatives were mostly enriched in the replicative senescence model (high passage). Monosaccharides, fatty acylcarnitines and dipeptides were commonly enriched also in irradiated samples, and cells treated with etoposide. In the hydroxyurea-treated cells, we detected an enrichment of pyrimidines, hydroxycinnamic and phenylpropanoic acids.

Of notice, in the media of the late passaged cells we detected high levels of prostaglandins (PGF1α, and PGF2α), and low levels of catecholamines (methyldopamine). This latter was also detected at low levels in cells treated with 12Gy IR and 800μM hydroxyurea, while in the 10μM etoposide-treated cells we observed high levels of methyl-dopamine.

Regarding the intracellularly detected metabolites, among the different senescence-induced models we identified common classes of amino acids, TCA acids, pyrimidine ribonucleotide diphosphates, amines, dicarboxylic acids, dipeptides and saturated fatty acids. Pyridoxamines were representative of cells at late passage (P=20) and the irradiated cells. Fatty acyl carnitine, purine ribonucleotide monophosphates and sugar acids were also measures in the hydroxyurea-treated cells. While unsaturated fatty acids and hippuric acids were represented in the late passaged cells and the ones treated with hydroxyurea. The classes of metabolites shared between irradiated and hydroxyurea-treated cells were amino alcohols and monosaccharides. Monosaccharides phosphates and imidazoyl carboxylic acid were also detected upon etoposide treatment. Pyrimidones were shown upon irradiation and etoposide senescence induction. Peculiar for each phenotype were: cyclohexylamines in cells at late passage, amino benzoic acids, and glycosylamines in cells treated with hydroxyurea, and peptides in the irradiated cells.

## Discussion

Identifying senescent cells for the treatment of age-related disease is a primary unresolved challenge in ageing research. Thus, the development of cell models to understand senescent phenotypes with the aim of developing biomarkers is of huge importance. Heterogeneity of the senescence phenotype represents a significant barrier to studying the mechanisms underpinning the phenotypes of ageing, the consequences of which include the lack of a universal model of senescence and a universal marker for the identification of senescent cells [29].

In this study we generated different models of senescence based on replication stress (replicative senescence), genotoxic and oxidative stress (stress-induced senescence) stimuli. The rationale behind the selection of these models was to explore how different stress stimuli influence the development of senescence through the accumulation of various biomolecular and metabolic alterations.

High ß-Gal expression, reduced proliferation, cell enlargement and flattening are primary characteristics of senescent cells. Initially we evaluated the presence of those phenotypic markers in cells at multiple passages and treated them with multiple dosages of irradiation, hydroxyurea or etoposide (**Figure 1**, **Error! Reference source not found. of Appendix 1**, **Error! Reference source not found. of Appendix 1**, **Figure 2**, **Figure 3**). From our results, cells at a late passage (P = 20) showed a higher level of ß-Gal expression, in line with other studies [8]. Ionizing radiation (IR)-induced senescence has been extensively studied showing induction by exposure to moderate IR doses [30]. Our study further confirms this evidence of senescence induction at 12Gy irradiation. Hydroxyurea induced senescence-like changes in normal human fibroblasts after long-term treatment with higher drug concentrations (400-800 μM) [31]. Conversely, etoposide is known to induce senescence at a low dosage (<50 μM) [32]. Accordingly, our results showed a global increase of ß-Gal expression at 800 μM of hydroxyurea and 10 μM of etoposide. At the morphological level, senescent cells present a characteristic enlarged and flattened morphology [33]. According to this, the human fibroblasts of this study at late passage (p=20), and at dosage of 12 Gy irradiation, 800 μM hydroxyurea and 10 μM etoposide, increased their length and width compared to the early passaged cells and cells treated at lower radiation and chemical dosage. Of notice, in the chemical treated cells higher concentrations of hydroxyurea (1000μM) and etoposide (25-50 μM) resulted in reduction of the cell size (width and length) which might be associated with the treatment toxicity and the induction of programmed cell death or apoptosis. Shrinkage of the cell is indeed a hallmark of apoptosis [34], however, specific molecular markers are needed to verify this assumption.

From analysis of γH2AX foci DSB immunolabelled foci (**Figure 2**), higher DNA damage foci were observed in drug-treated cells (hydroxyurea and etoposide) compared to cells undergoing replication-induced senescence and treatment with IR. While the analysis of the Ki-67 replication marker showed a significant reduction of Ki-67-positive cells in the late passaged and etoposide treated cells (**Figure 3**). This reduction was not significant in cells exposed to IR or absent in hydroxyurea-treated cells. Based on these results we observed that accumulation of DNA damage and reduction of proliferation index are common traits of the stress-induced phenotypes of this study, therefore confirming their effective senescence-like phenotype.

Genetic and molecular biomarkers – including proteins – have been extensively used for the identification of senescent cells both in vitro and in vivo. They have been useful in underlying key biological features of senescence and identifying its major drivers during natural ageing and in age-related diseases. However, genetic and molecular biomarkers are generally nonspecific and non-reproducible across all cell/tissue types [35]. These limitations represent an obstacle for the effective detection of senescence and a proper understanding of its mechanisms and role in living organisms.

Metabolic reprogramming is another hallmark of senescent cells and adaptive response to maintain their viability in a growth-arrested state [17]. Metabolic changes are highly susceptible to the type of stress stimuli, contributing to the heterogeneity of the senescence phenotypes. On this basis, metabolomics has recently emerged in ageing research as a fundamental tool to depict the complexity of senescence, ageing, and the correlation with the accumulation of damages [36].

Our goal in the present study was to apply a combined analysis of senescence molecular biomarkers with global untargeted mass-spectrometry-based metabolomics to map the metabolic changes occurring in response to different senescence-inducing stimuli. We examined the metabolic differences across the different models of senescence including replicative senescence and SIPS. A general enrichment of intracellular metabolites was computed for the senescence model induced by replication, irradiation and hydroxyurea stress, while depletion of metabolites was observed upon etoposide treatment (**Figure 4**). We found overlapping intracellular metabolic pathways (histidine, alanine, aspartate, glutamate, and arginine metabolism) across the four senescent models (**Figure 5**, **Error! Reference source not found. of Appendix 1**). We then investigated intracellular metabolic pathways that were unique to specific senescence phenotypes (Irradiation, late passaged and etoposide: glutamine and glutamate metabolism; hydroxyurea and late passaged: arginine and proline metabolism; etoposide: glycine, serine and threonine metabolism).

Regarding the extracellular metabolome, we detected a general trend towards the enrichment of metabolites in all the senescence-induced models including etoposide-treated cells **(Figure 8)**. The analysis of the class of metabolites that were mainly affected by metabolic changes upon induction of senescence, revealed that amino acids were the most susceptible to these changes both in the intracellular and extracellular metabolome of all the senescence-induced cells (**Figure 9**). Below, we will discuss the level of the most relevant metabolic changes affecting amino acids and some of the derived metabolic pathways (**Figure 7**), and the role they might play in the context of senescence associated pathogenesis.

Regarding histidine metabolism from the analysis of intracellular metabolites in the senescence-induced phenotypes, we observed a general increase of the histidine-derived catabolic product, methylhistidine, indicative of enhanced proteolysis of histidine which is a marker of frailty [37]. Depletion of ergothioneine, another histidine-derived metabolite, was observed in all senescence-like phenotypes. Ergothioneine is a powerful scavenger of hydroxyl radicals, thus it plays an important role against oxidative damage [38]. Low levels of ergothioneine have been found in patients with Parkinson’s disease [39] and the elderly [40].

Accumulation of phenylalanine was observed in the replicative senescence model in line with the study of James EL., at al 2015 [41]. However, in the cells exposed to IR and chemical-treated cells phenylalanine was depleted, which might be attributed to its higher consumption. Elevated serum phenylalanine has been linked to telomere loss in men [42], inflammatory diseases [43], and type 2 diabetes [44]. Of notice, phenylalanine is a precursor for the synthesis of catecholamines including tyramine, dopamine, epinephrine and norepinephrine. High levels of nitrotyramine were observed in the media of hydroxyurea cells, and high levels of methyldopamine and norepinephrine have been observed in the media of etoposide cells.

Catecholamines are hormones involved in the immune response [45] with roles in cellular proliferation and apoptosis [46]. Catecholamines act as anti-inflammatory molecules decreasing the levels of TNFα, CCL2, IL-6 and IFNα [47]. This is suggesting that alanine metabolism in certain senescence-like phenotypes can play a significant role in the inflammatory response of these cells.

In agreement with previous studies we observed increased arginine biosynthesis through the urea cycle across all senescence conditions, except for etoposide-treated cells [41]. Accordingly, high levels of urea, a waste product of arginine metabolism, have been detected in all the cell lines including etoposide. Proline is derived from the urea cycle and low levels of this metabolite have been detected across all the senescence-induced models in this study. Proline is a metal chelator that supports oxidative damage [48] and can be hydroxylated to form hydroxyproline whose levels have been found to be decreased in the media of all the senescence-induced cells. Diminished hydroxylation of proline is diminished under hypoxia conditions which are known to induce cell cycle arrest [49], although its correlation with senescence is still debated [50].

Increased glutaminolysis due to overexpression of GLS1 has been observed in previous studies [51]. High levels of glutamine were observed at later passages, together with high levels of glutamate, α-KG, malate and aspartate which are intermediate of the TCA cycle. This suggests a role for glutamine in sustaining the TCA cycle activity in replicative senescent cells.

Previous work has found that ATP is generally depleted in senescent cells [41]. However, in the present work ATP was not significantly enriched in response to the induction of senescence through replication stress. Further analyses are needed to investigate the mechanisms of energy metabolism in HFF-1 cells after multiple passages in comparison with other cell lines. On the other hand, the SIPS models of senescence showed accumulation of AMP and a general downregulation of TCA metabolites.

Elevated levels of D-serine were observed in all the senescent phenotypes induced in the present work, while L-serine was not detected in any of the induced phenotypes. Depletion of some L-amino acids has been associated with the induction of stress signals by activation of the general control non-depressible 2 (GCN2) [52], responsible for the phosphorylation eIF2a and the consequent production of pro-inflammatory cytokines [53, 54]. Accordingly, significantly elevated levels of prostaglandins F1a and F2a were detected in the media of the replicative senescence model. Moreover, the intracellular level of prostaglandins derived from the metabolism of arachidonic acid were higher in these cells. Depletion of serine-derived cystathionine was also observed for all the conditions accompanied by reduction of taurine which is a product of cysteine metabolism [55]. Several studies associated reduced levels of taurine to senescence [56]and age-related diseases [57, 58] where taurine plays as an antioxidant and inhibitor of pro-inflammatory molecules [59].

A targeted metabolomics analysis of these senescence-induced phenotypes is required to validate our findings. This could allow the further validation and development of metabolic markers of senescence, informing future studies into the mechanism underpinning senescence induction in response to different stimuli. In medicine, the identification of new and specific biomarkers of senescence will be beneficial in the measurement of senescence burden within an individual to prevent age-related dysfunction and diseases, and predict response to treatment/surgery. Encouraging studies and clinical trials are showing the therapeutic effects of the pharmacological elimination of senescent cells through compounds called senolytics (inducers of senescent cell death) and senomorphics (SASP inhibitors) [60]. Hence, biomarkers of senescence will also help to assess the dosing of senolytic/senomorphic drugs in the clinical assessment of senescence burden specific for each individual.

Moreover, a limitation of the present work was analysis of senescence induction in a single cell type. Future studies require the assessment of cellular response to senescence induction in a panel of cell lines for a more comprehensive analysis of various senescence phenotypes and their manifestation in cell lines originating from different tissue types. Understanding the mechanisms of senescence induction in different tissue types will aid in cataloguing the specific intra- and extracellular features associated with cellular senescence.

## Conclusion

Our results have shown intra- and extracellular metabolomics profiles of different models of senescence including replicative and stress-induced senescence through DNA damage and ROS induction. We presented metabolomics as a powerful strategy for the identification of human cellular senescence biomarkers which characterise a wide variety of human pathologies including ageing, frailty, inflammation, cardiovascular, neurodegenerative diseases, and cancer. Moreover, these findings may provide potential therapeutic targets for preventing the development of age-related diseases, or as adjuvant therapies to improve patients’ response to treatments.

## Materials

### Cell line, chemical and treatment

All cell culture reagents were obtained from Gibco (Thermo Fisher Scientific). Human Foreskin Fibroblasts cell line (HFF-1, ATCC^®^ SCRC-1041) was purchased and maintained in Dulbecco’s Modified Eagle’s Medium (DMEM, high glucose) supplemented with 10% *v/v* FBS (high glucose, Invitrogen), 1% *v/v* non-essential amino acids (NEAA), and 1% *v/v* penicillin-streptomycin (Invitrogen). Cells were maintained in a pre-humidified atmosphere containing 5% *v/v* CO_2_ at 37°C.

Hydroxyurea (Sigma Fisher Scientific) was prepared as a 10 mM stock in water. Etoposide was prepared as a 1 mM stock solution in DMSO. All drug stocks were aliquoted and stored at - 20°C until use.

Crystal violet (Sigma Fisher Scientific) stain was prepared with 20% methanol (Alfa Aesar by Fisher Scientific) and 2% sucrose (VWR Life science) and stored at ambient temperature. Primary antibodies for γH2AX and Ki67 (Cell Signalling Technologies) were used for foci immunostaining alongside Alexa Fluor^®^ 488-conjugated secondary antibody (Fisher Scientific).

Cells were passaged multiple times in culture (up to 20 passages) and were either treated with ascending doses of irradiation (1-12 Gy for 1 week), hydroxyurea (0-1,000 μM for 2 weeks), or etoposide (0-50 μM for 1 week) until markers of senescence appeared (growth arrest, increase in cell size, ß-Gal expression).

## Methods

### Senescence associated ß-galactosidase staining

Cells were seeded in a 96-well plate at a density of 4,000 per well and incubated for a pre-defined period for each specific treatment type. Following incubation, senescence associate ß-Gal activity was assessed using a senescence detection kit (ab65351, Abcam) as per the manufacturer’s recommendations. Subsequently cells were counterstained with DAPI to stain their nuclei. Image acquisition was carried out using an Invitrogen EVOS Auto Imaging System (AMAFD1000-Thermo Fisher Scientific) with a minimum of 100 cells imaged per treatment condition. ß-Gal-stained cells were manually counted with ImageJ software. The number of ß-Gal-stained cells was normalized with the number of counted nuclei.

### Cell viability assay

The crystal violet assay was used to determine the viability of human fibroblasts following the induction of senescence. HFF-1 cells were plated in 96 well plates at 4,000 cells per well, treated and incubated for pre-defined times depending on the mechanism of senescence. Following treatment, the culture medium was removed, and the treated cells were washed once with PBS and stained with the crystal violet staining solution for 30 min at ambient temperature. Subsequently, the staining solution was removed, cells were gently rinsed with tap water and left to dry for 24 hours at ambient temperature. Subsequently, the stain was solubilized with 100% Ethanol and quantified at 600 nm using a GM3500 Glomax^®^ Explorer Multimode Microplate Reader (Promega).

### Immunostaining for γH2AX and Ki67

Foci immunodetection for γH2AX and Ki67 was performed in low and high passage cells seven days after seeding for non-irradiated cells and for cells irradiated at 12Gy for seven days, in non-treated and treated cells with hydroxyurea (800 μM) and etoposide (10 μM) for 14 and seven days, respectively. Briefly, cell monolayers were fixed in chilled 4% w/v formaldehyde containing 2% w/v sucrose in PBS, followed by fixation in ice-cold methanol (100% v/v). Subsequently, cells were permeabilized in 0.25% v/v Triton X-100 in PBS, blocked with 5% v/v goat serum/5% w/v BSA, immunoprobed with either a primary rabbit anti-γH2AX (1:1000) or primary mouse anti-Ki67 (1:1000) antibody overnight at 4°C. Cell monolayers were treated with goat, anti-rabbit and anti-mouse Alexa Fluor^®^ 488 conjugated secondary antibody and counterstained with DAPI. Image acquisition was carried out using an Invitrogen EVOS Auto Imaging System (AMAFD1000-Thermo Fisher Scientific) with a minimum of 100 cells imaged per treatment condition. Resultant γH2AX foci and Ki-67 labelled nuclei images were analysed in Cell Profiler (v.4.2.1.) using a modified version of the “speckle counting” and “percent positive” pipeline, respectively. Further thresholding setting for nuclei, γH2AX foci and Ki-67 labelled nuclei are indicated in **Table 1**.

**Table 1.** Thresholding parameters applied in Cell Profiler for the detection of foci and labelled nuclei.

|  | **DAPI** | **γH2AX** | **Ki-67** |
| --- | --- | --- | --- |
| Diameter of objects (in pixel units) | Min: 8<br>Max:80 | Min: 1<br>Max: 40 | Min: 5<br>Max:200 |
| Distinguish objects by | Shape | Intensity | Intensity |
| Thresholding method | Minimum<br>Cross-Entropy | Minimum<br>Cross-Entropy | Minimum<br>Cross-Entropy |
| Lower and upper bounds on threshold | 0.05, 1.00 | 0.00, 1.00 | 0.00, 1.00 |

### Sample preparation and metabolite extraction

HFF-1 cells were seeded at a density of 2×10^6^ cells per well in 6-well plates, at passage 20 and before exposure to either 12 Gy irradiation, or growth medium containing 800 μM hydroxyurea or 10 μM etoposide. Following senescence induction, the growth medium was aspirated from each well, centrifuged to remove cell debris, aliquoted and stored at −80 °C. Next, treated cells were washed with pre-chilled PBS, with the metabolites quenched and extracted in a final volume of 1.5ml pre-chilled (−80°C) methanol:acetonitrile:water solvent (50:30:20). Resultant cell pellets were collected, and flash frozen in liquid nitrogen, vortexed and sonicated for three minutes in an ice-water bath. This process was performed in triplicate. Resultant extracts were centrifuged at 13,000xg for 10 minutes at 4°C and the pellets retained for protein quantification using the Bradford assay (Pierce™ Coomassie Plus Bradford assay kit, Thermo Scientific™). The resultant supernatant was collected, and dried with a Speed vac centrifuge for 10 h (Savant-SPD121P). Dried metabolite pellets were subsequently reconstituted in acetonitrile:water (50:50) at volumes normalised to the relative protein content. Quality control (QC) samples were prepared by pooling samples across all control and treatment groups. Solvent blank (reconstitution buffer) and QC samples were inserted in the analytical batch.

### Liquid Chromatography Tandem Mass Spectrometry (LC-MS/MS)

Metabolite separation was performed on a binary Thermo Vanquish ultra-high-performance liquid chromatography system where 10 μl of the reconstituted cellular extract was injected onto a Thermo Accucore HILIC column (100mm x 2.1mm, particle size 2.6μm). Blank and QC samples were injected after every five samples to assess the stability of the detecting system. The temperature of the column oven was maintained at 35°C while the autosampler temperature was set at 5°C. For chromatographic separation, a consistent flow rate of 500μl/min was used where the mobile phase in positive and negative heated electrospray ionisation mode (HESI+/-) was composed of buffer A (100% Acetonitrile with 0.1% formic acid) and buffer B (20 mM ammonium Acetate in water with 0.1% formic acid).

A high-resolution Exploris 240-Orbitrap mass spectrometer (Thermo Fisher Scientific) was used to perform a full scan and fragmentation analyses. Global operating parameters were set as follows: spray voltages of 3900V in HESI +ve mode, and 2500V in HESI −ve mode. The temperature of the transfer tube was set at 320°C with a vaporiser temperature of 300°C. Sheath, aux gas and sheath gas flow rates were set at 40, 10 and 1 Arb, respectively. A top-5 Data-dependent acquisition (DDA) was performed using the following parameter: survey scan range was 50-750 m/z with MS1 resolution of 60,000. Subsequent MS/MS scans were processed with a resolution of 15,000. High-purity nitrogen was used as nebulising and as the collision gas for higher energy collisional dissociation.

### Mass Spectrometry Data Processing

Raw data files obtained from Thermo Scientific Xcalibur TM software v4.2 were imported into Compound Discoverer™ 3.3 software where the “Untargeted Metabolomics with Statistics Detect Unknowns with ID Using Online Databases and mzLogic” feature was selected. The workflow analysis performed retention time alignment, unknown compound detection, predicted elemental compositions for all compounds, hid chemical background (using Blank samples). For the detection of compounds, mass and retention time (RT) tolerance were set to 3 ppm and 0.3 min, respectively. The library search was conducted against the mzCloud, Human Metabolome Database (HMDB) and Chemical Entities of Biological Interest (ChEBI) database. A compound table was generated with a list of putative metabolites (known and unknown). Among them, we selected all the known compounds fully matching at least two of the annotation sources. The selected metabolites were then used to perform pathway and statistical analysis.

### Pathway Analysis with MetaboAnalyst

Prior to the analysis of the intracellular and extracellular metabolic pathways with MetaboAnalyst 5.0 (http://www.metaboanalyst.ca/), a HMDB identification code was assigned to each selected metabolite. For intracellular metabolites, joint pathway analysis was performed by mapping the altered metabolic pathways onto senescence-associated genes with the list of ID compounds and their associated Log2 Fold change values. The integration method combined both genes and metabolites into a single query, which was then used to perform the enrichment analysis based on a hypergeometric test. The hypergeometric test allows to define the significance of the association between two sets of data (genes and metabolites) [61]. Finally, important nodes (compounds) were scored based on their betweenness centrality, and pathway analysis results were generated.

For the extracellular metabolites Metabolite Set Enrichment Analysis (MSEA) was applied by uploading a list of ID metabolites entered as one-column data followed by a hypergeometric test to evaluate whether a particular metabolite set is represented more than expected by chance within the given compound list.

### Statistical analysis

All data are presented as mean ± standard deviation (n≥5). For metabolomics analysis, Principal Component Analysis (PCA) was performed to test the analytical reproducibility of QC injections, reduce the dimensionality of data, and visualise the presence of any clustering differences between sample groups. Differential analysis was used to compare differences between control and treatment groups and plotted as a Volcano plot (log2-fold change vs. - log10 p-value). Peak areas were log10 transformed and p values calculated for the sample group by analysis of variance (ANOVA) test. A p value<0.05 and a fold-change >1.5 was deemed to be statistically significant.

